# Mechanoresilience of lysosomes conferred by TMEM63A

**DOI:** 10.64898/2025.12.18.695245

**Authors:** Angelina S. Kim, Jing Ze Wu, Ruiqi Cai, Spencer A. Freeman

## Abstract

Lysosomes are subject to perturbations that can cause damage to their limiting membrane. Osmotic shifts, pore-forming toxins, and the growth of luminal polymers or pathogens all stand to increase lysosomal membrane tension and/or disrupt the bilayer. In some contexts, this leads to lysosomal rupture and cell death. Here we describe a mechanism that enables lysosomes to sense and respond to acute increases in tension of their limiting membrane. We report that the lysosome-resident non-selective cation channel, TMEM63A, can drive the directional flux of monovalent cations, major osmoticants, out of the lumen when gated by mechanical tension on the organelle. This results in the ability for lysosomes to relieve hydrostatic pressure and, proportionally, membrane tension, affording lysosomes the time to acquire additional lipids. Lysosomes without this mechanism −either because TMEM63A is deleted or in the case when cells express disease-causing variants of *TMEM63A*− are an order of magnitude more sensitive to lysis upon increases to their membrane tension when compared to their wildtype counterparts. These findings suggest that lysosomes are capable of regulating hydrostatic pressure and volume in response to high tension.

## Introduction

Lysosomal damage contributes to a wide variety of pathophysiological settings. These range from infection, where endocytosed pathogens grow or release effectors that perforate endomembranes (Gaillard et al., 1987), to neurodegenerative disease, which can be associated with the aberrant aggregation of luminal fibrils in lysosomes (Filimonenko et al., 2007). In turn, the damage to lysosomes leads to tissue wide responses, notably inflammation (Duewell et al., 2010). If left unchecked, lysosomal damage causes cell death (Hornung et al., 2008). Accordingly, there is an emerging appreciation for mechanisms involved in sensing lysosomal stress or damage and repairing the injured membrane.

Many of these responses centre on increases to cytosolic calcium, proposed to leach from compromised organelles. The local gain in cytosolic calcium can conceivably lead to lysosome repair in a number of ways. Cytosolic calcium activates scramblases on the lysosome, exposing sphingomyelin to neutral sphingomyelinase (Niekamp et al., 2022). The cleavage of sphingomyelin to generate conically shaped ceramide imbues the inward curvature required for intraluminal vesiculation, an effect that stands to sort damaged or damaging components inward. Calcium also recruits and activates ESCRT complexes to the lysosome that remodel the membrane and provide resilience to damage (Chen et al., 2024; Skowyra et al., 2018). Finally, calcium could support fusogenic complexes to stimulate the recruitment of additional membrane via fusion with other (endo)lysosomes (Westman et al., 2020). This latter effect is critical as membrane bilayers are minimally distensible, stretching only 2-5% before tension exceeds what the lipid packing can tolerate before undergoing rupture (Evans et al., 1976; Koslov and Markin, 1984). An additional set of secondary response pathways involve the directional flow of lipids to tensed lysosomes from the ER across membrane contact sites. Both VPS13C and PDZD8 have been proposed to achieve these ends and are important for the comparatively slow expansion of the delimiting lysosomal membrane upon stress (Durgan and Florey, 2022; Wang et al., 2025; Yang et al., 2025).

An important question emerging from the preceding observations is what are the mechanisms that *directly* sense lysosomal stress to initiate the responses that prevent rupture? Even in the absence of infection or injury, lysosomes are subjected to threats to their integrity yet are not damaged. To date, the sensory machinery that provide these degradative organelles with resilience against acute rupture are unknown. The recent discovery that members of the OSCA/TMEM63 family of mechanosensitive cation channels localize to vacuoles/lysosomes in cells suggests the possibility that these channels could be direct sensors of lysosomal stress (Li et al., 2024; Murthy et al., 2018). By outwardly conducting cations upon mechanical gating, OSCA/TMEM63 could also support both osmotic and/or calcium-dependent aspects of the response.

By acutely increasing the hydrostatic pressure and membrane tension on lysosomes, we find that these organelles withstand damage by the activation of TMEM63A. TMEM63A relieves high membrane tension by rapidly alleviating osmotic pressure. This enables the secondary responses that deliver additional membrane surface to lysosomes either through homotypic fusion or the transport of lipids from the ER, leading to the lysosome enlargement that protects from rupture. Deletion of TMEM63A leaves cells highly vulnerable to lysosomal cell death. These results demonstrate that organelles are capable of regulated tension and volume decrease as a primary mechanism of resilience in response to stress.

## Results

### Lysosomes withstand high membrane tension

Numerous experimental conditions have been established that lead to lysosome stress and rupture. We performed dose-response or time-course experiments in primary bone-marrow derived macrophages (BMDM) and RAW264.7 macrophages with the cell-permeant lysosomotropic agents, L-Leucyl-L-Leucine methyl ester (LLOMe) or Glycyl-L-phenylalanine 2naphthylamide (GPN), as well as sucrose or alum, which slowly accumulate in lysosomes by fluid phase endocytosis. If the limiting membrane of the lysosome undergoes rupture, it leads to the recruitment of galectin scaffolds that detect the exposed glycans that comprise the luminal lysosomal glycocalyx (**Fig 1A**) which is easily scored. Additionally, membrane rupture causes the release of its luminal contents including small membrane-impermeant dyes loaded by endocytosis, e.g. sulforhodamine G (SRG), and alkalinization, which can be assessed with Lysotracker (**Fig 1A**). With these approaches, we determined the concentrations required for LLOMe or GPN to cause rupture of the lysosomes in macrophages (**Fig 1B-I**). In contrast, we did not see rupture of lysosomes in the cells at any time after their incubation with sucrose or alum (**Fig 1J-L, S Fig 1A-C**).

**Figure 1.**
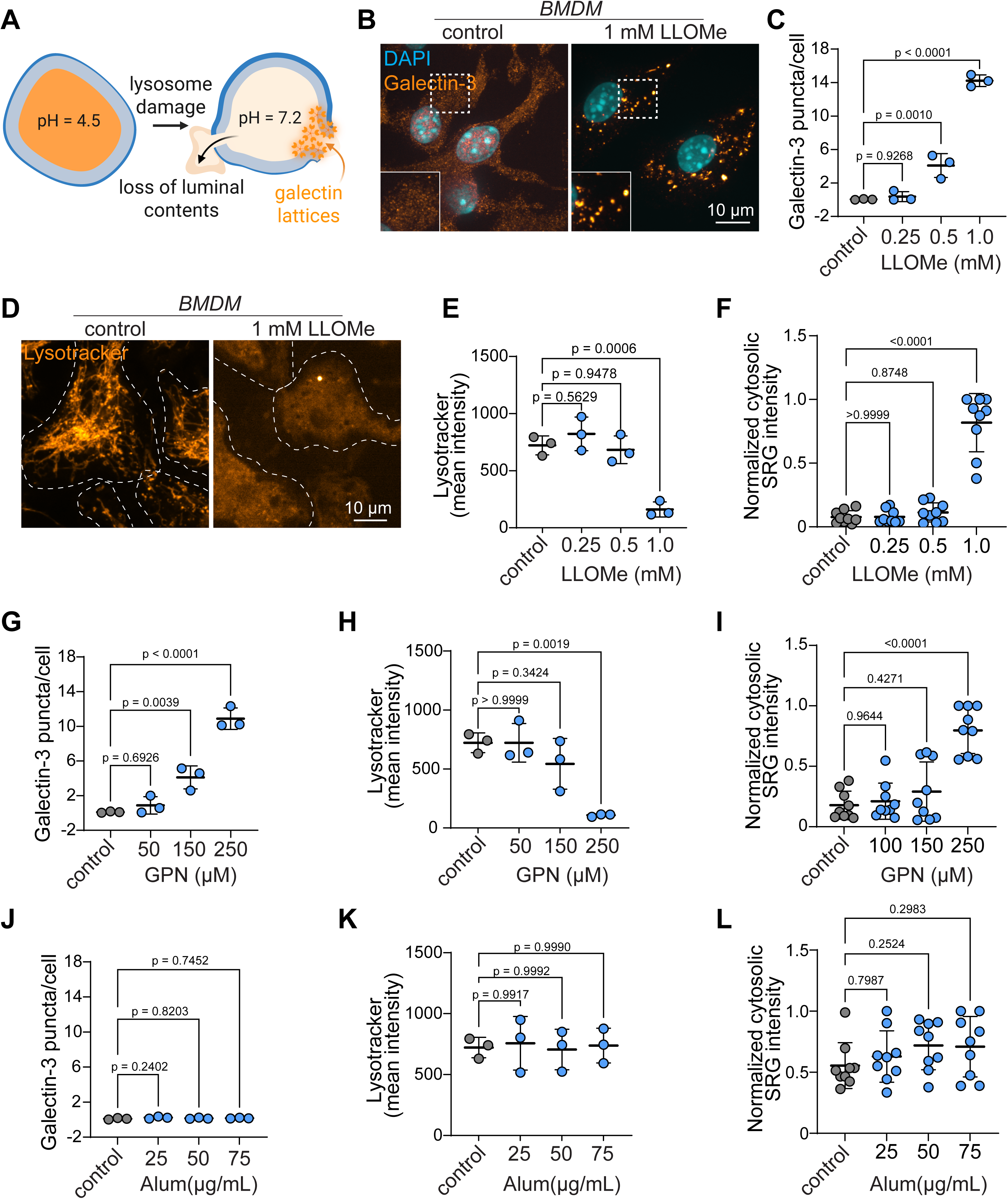
Lysosome damage and resilience in macrophages. **A)** Model. **B-F)** Lysosomal damage upon treating WT primary macrophages with indicated concentrations of LLOMe for 10 min was determined by staining for galectin-3 puncta **(B-C)**, measuring the loss of lysosome acidification with Lysotracker **(D-E)**, or quantifying cytosolic sulforhodamine G (SRG) signal **(F)**. The same was done for GPN for 10 min **(G-I)**, and Alum for 3 h **(J-L)**. Each point for galectin-3 puncta and Lysotracker quantification represents mean of experiment (>5 cells per field, 15 fields in total). Each dot for SRG represents a field (>5 cells per field, 9 fields in total). n = 3 independent experiments, scale bars 10 µm.

We went on to determine if tension is increased before rupture in the LLOMe or GPN experiments, which we have previously characterized for sucrose-laden lysosomes. This was done using the recently described Lyso-Flipper probe (Colom et al., 2018). Briefly, cells were treated with concentrations of LLOMe or GPN for time periods that did not cause lysis of the lysosomes. Under these settings, we noted increased lifetime of the Lyso-Flipper probe, suggesting the limiting membrane incurs increases in its lateral tension upon such perturbations (**Fig 2A**). An osmotic gain in by the membrane-impermeant products (glycyl-L-phenylalanine and 2-naphthylamine) of GPN produced upon its cleavage by cathepsins is expected to generate proportional increases in membrane tension. Like GPN, LLOMe is cleaved by lysosomal cathepsins, but this results in a hydrophobic (Leu-Leu)n-OMe polymer (Thiele and Lipsky, 1990); the increased tension upon LLOMe may either result from pressure produced by the polymers directly or their membrane disruptive properties.

**Figure 2.**
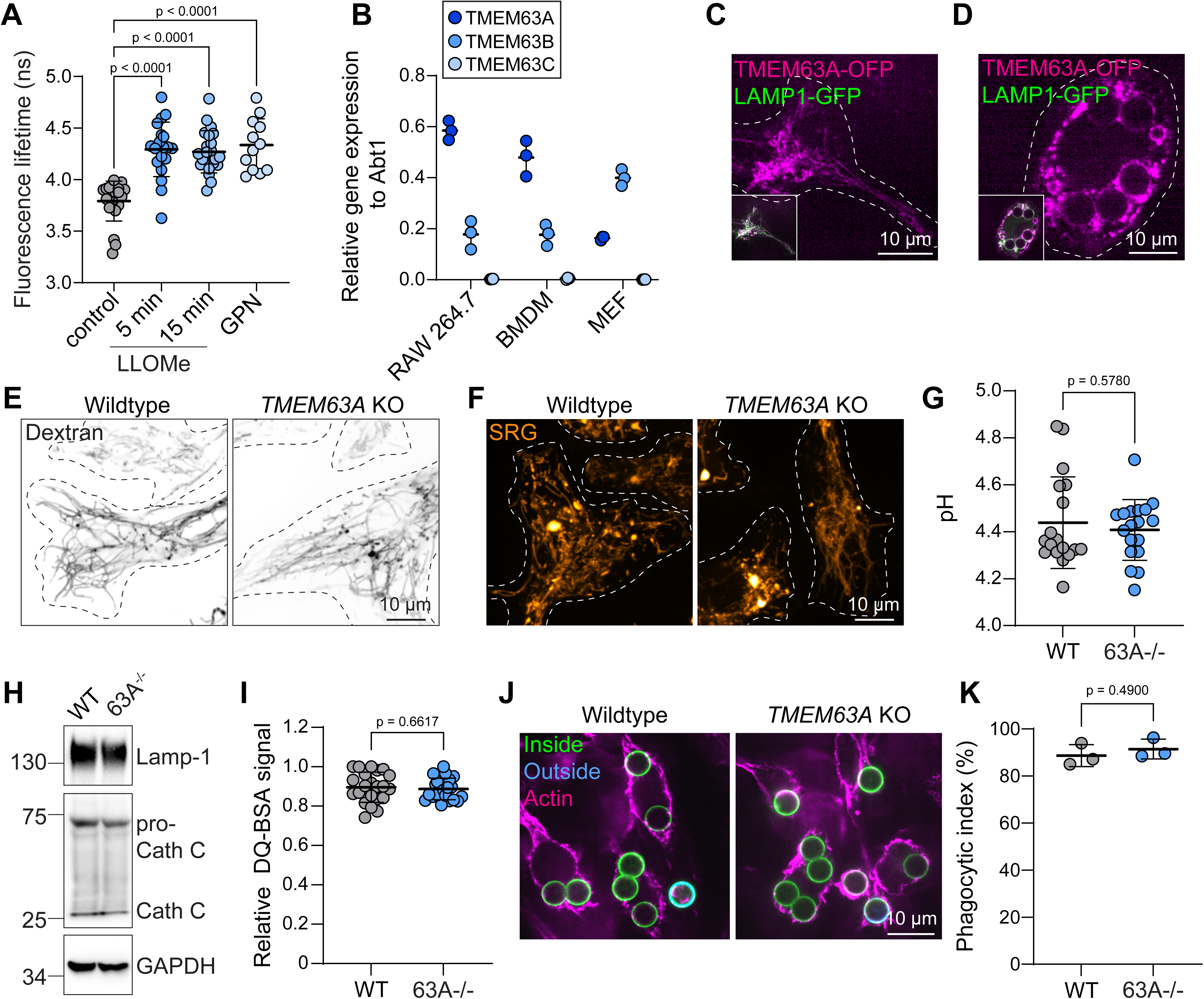
Lysosomal membrane tension and TMEM63A expression and targeting in macrophages. **A)** LysoFlipper FLIM in WT RAW264.7 macrophages incubated with 1 mM LLOMe for 5 or 15 min, or 250 µM GPN for 15 min. Each dot represents one field of 5-10 cells. N = 3. **B)** qPCR for TMEM63A, B, and C in RAW264.7, BMDM, or MEF. n = 3. **C-D)** TMEM63A-OFP (magenta) and LAMP1-GFP (green) in WT RAW264.7 cells before and after challenge with IgG-coated microspheres. **E-F)** 10 kDa dextran or SRG pulsed for 16 h and chased for 1 h in WT and TMEM63A KO BMDM. **G)** Lysosomal pH in WT and TMEM63A KO BMDM determined using ratiometric measurements of 10 kDa Oregon Green dextran pulsed for 16 h and chased for 1 h. Each data point represents a field containing >5 cells. n = 3. **H)** LAMP1, cathepsin C expression and processing by WB. **I)** DQ-BSA signal of WT and TMEM63A KO BMDM after pulse (1 h) and chase (1 h). All data points represent a field containing 4-8 cells. n = 3. **J-K)** Phagocytosis in WT and TMEM63A KO BMDM, quantified in K. Each data point represents 3-5 fields containing 15-25 cells. n = 3, scale bars 10 µm.

### TMEM63A localizes to lysosomes

Under such conditions, mechanosensitive channels could be opened. To determine which channels would be positioned to respond to increases in lysosome membrane tension, we first analyzed the expression of TMEM63A, B, and C in macrophages. TMEM63A and B were both well-expressed in macrophages; TMEM63A was expressed at 3- and 10-fold higher levels in macrophages compared to murine fibroblasts (**Fig 2B**). We found that TMEM63A showed pronounced localization to lysosomes and phagosomes as indicated by coincidence with LAMP1-GFP (**Fig 2C-D**) (Li et al., 2024) suggesting that TMEM63A may play a role in the lysosomes of highly endocytic cell types.

### The loss of TMEM63A does not affect lysosome morphology, pH, or cause susceptibility to damage in response to the slow accumulation of solutes and particles

To test for the role of TMEM63A in lysosomes of macrophages, we generated TMEM63A KO cells and TMEM63Afloxed LysM-Cre conditional KO mice (**S Fig 1D-F**). We initially investigated the morphology of the tubular lysosome network in the macrophages derived from the WT versus conditional KO mice. The morphology of lysosomes in the TMEM63A KO cells appeared to be normal and these retained 10 kDa dextran and SRG (**Fig 2E-F**). We also noted no differences in lysosomal pH in the TMEM63A KO cells (**Fig 2G**) or differences in LAMP1 or cathepsin C expression (**Fig 2H**). DQ-BSA was also taken up and degraded normally in the TMEM63A KO cells (**Fig 2I**) and the cells were capable of phagocytosis, even when challenged with a large number of phagocytic targets (**Fig 2J-K**). Taken together, these data suggested that, under steady-state conditions, there were no obvious defects in the lysosomes in the TMEM63A KO macrophages. We therefore proceeded to determine the function of TMEM63A under conditions in which it would presumably become activated, i.e. under high tension.

Tension on the limiting membrane of lysosomes increases upon their slow accumulation of solutes including sucrose which cannot be digested nor outwardly transported. The same occurs with the vaccine adjuvant alum. We found that neither insult caused the overt rupture of lysosomes in WT macrophages despite causing their growth, ostensibly by fusion (**Fig 3A-E**). We also did not find that the lysosomes in TMEM63A KO cells underwent any signs of rupture upon accumulating sucrose or alum (**Fig 3A-E, S Fig 2C-D**) or upon phagocytosis of latex microspheres (not shown), suggesting that the channel is not required to preserve (phago)lysosomal integrity when solutes/particles slowly accumulate or during phagocytosis itself. These data also suggest that TMEM63A is not required for maturation in the endocytic pathway or the fusion of lysosomes.

**Figure 3.**
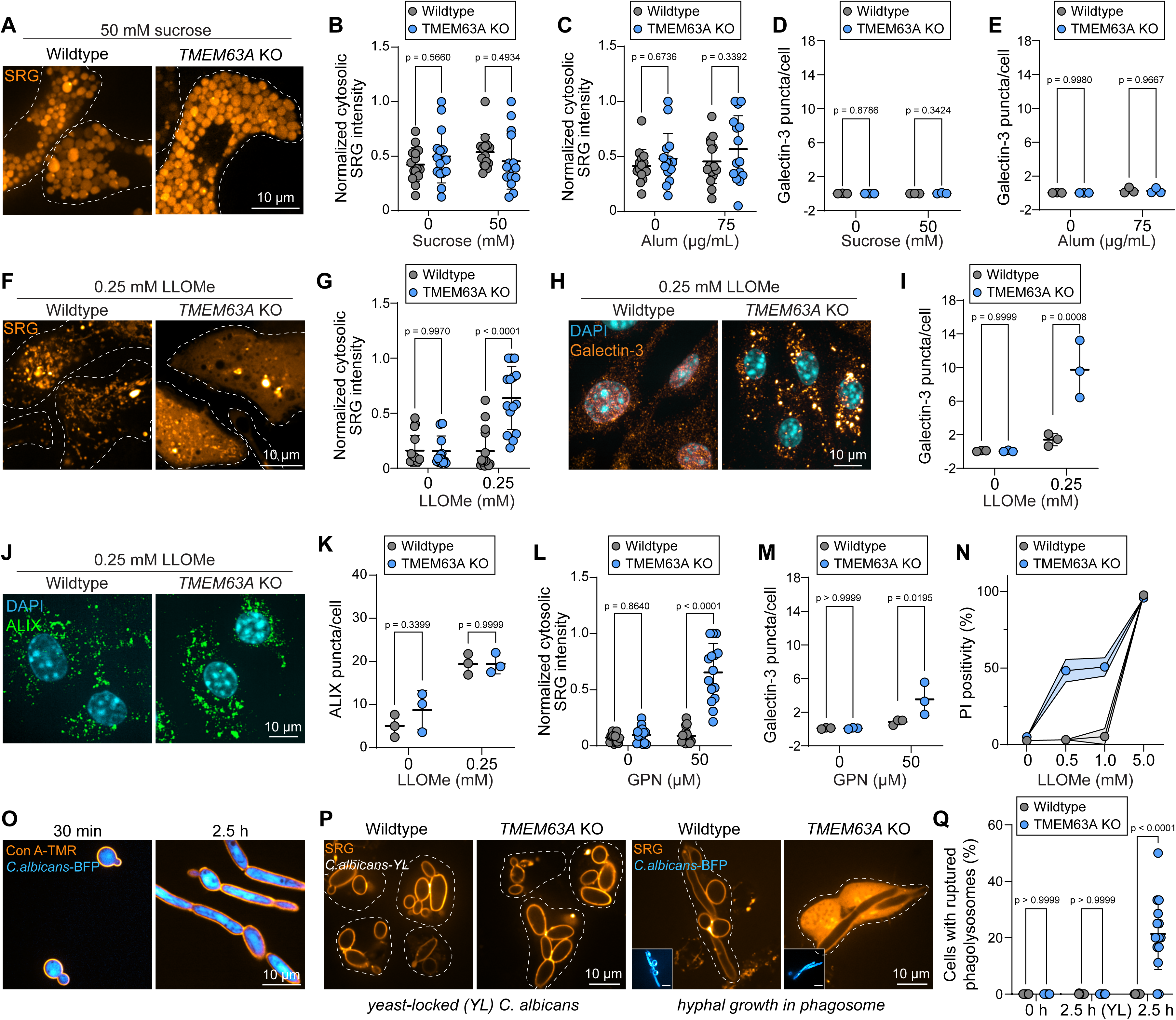
TMEM63A protects cells from lysosomal damage and cell death. A-E) WT or TMEM63A KO BMDM given SRG and either 50 mM sucrose overnight or 75 µg/mL alum for 4 h. Cytosolic SRG signal **(B-C)** quantified per field of 5-10 cells or galectin-3 puncta per cell **(D-E)** each data point representing 5 fields. n = 3. **F-K)** WT and TMEM63A KO BMDM treated with LLOMe as in Figure 1. Lysosomal damage was quantified accordingly. In **J-K**, ALIX was immunostained and puncta above a threshold size per cell quantified. Each dot represents mean of experiment (>5 cells per field, 15 fields total), n=3. **L-M)** Damage in WT and TMEM63A KO BMDM upon GPN. n = 3. **N)** Cell death upon LLOMe treatment for 30 min in WT and TMEM63A KO RAW 264.7 cells was determined using propidium iodide. n = 3. **O-Q)** BMDM challenged with Δ*ece1 C. albicans*-BFP or yeast-locked (YL) *C. albicans*, internalized by phagocytosis and allowed to grow in the phagolysosome for 2.5 h (O). % of cells with ruptured phagosomes quantified per field of 4-8 cells in Q. n = 3, scale bars 10 µm.

### TMEM63A protects lysosomes from damage in response to acute swelling and lysosomotropic agents

In stark contrast to the stress accumulated on lysosomes when they cannot digest particles or transport solutes, sudden shifts in hydrostatic pressure can cause swift increases in lysosome tension. At high enough concentrations, agents like GPN and LLOMe cause swelling and rupture of lysosomes in minutes. We therefore challenged the TMEM63A KO macrophages with GPN and LLOMe across a range of concentrations established previously (*see Figure 1*). Under these conditions, we determined lysosome rupture by the loss of luminal contents (SRG) and the accumulation of galectin-3, indicative of membrane lysis. Remarkably, the lysosomes in TMEM63A KO macrophages underwent rupture at concentrations of lysosomotropic agents far below those required to rupture lysosomes in cells with the channel expressed (**Fig 3F-I, L-M, S Fig 2A-B**). While the re-repression of the wildtype version of human TMEM63A rescued the TMEM63A KO cells from lysosomal damage, as measured by galectin-3 recruitment, expression of disease-causing variants of TMEM63A, i.e. those causing hypomyelinating leukodystrophies (Yan et al., 2019), did not (**S Fig 3E**).

Upon investigating the recruitment of ESCRT, we found that the ESCRT-associated protein ALIX was recruited to an equal extent in TMEM63A KO cells upon lower doses of LLOMe as in WT cells (**Fig 3J-K**), indicating that a failure to recruit ESCRT was not ostensibly the cause of increased rupture. Indeed, a majority of the ALIX recruitment to stressed lysosomes in WT cells occurred without their rupture, as judged by galectin-3 (**S Fig 2F**), which is consistent with its role in protection. The recruitment of ALIX in TMEM63A KO cells could occur only after damage **(S Fig 2F),** perhaps not excluding the possibility that TMEM63A activation recruits ESCRT.

### Damage to lysosomes in TMEM63A KO cells causes lysosomal cell death

Lysosome-mediated cell death is a consequence of permeabilization of the limiting lysosomal membrane. Given the preceding observations, we tested if the TMEM63A KO macrophages were more susceptible to death upon treatment with titrated doses of LLOMe. Based on PI positivity, we found that the TMEM63A KO macrophages underwent cell death at much lower concentrations of LLOMe (**Fig 3N**), consistent with the findings of the increased lysosome damage sensitivity.

### TMEM63A protects from pathogen growth-induced phagolysosomal damage

Pathogens, including the opportunistic fungal pathogen *Candida albicans*, can grow within phagolysosomes by shifting to a hyphal growth state (**Fig 3O**). We challenged primary macrophages with two different strains of *C. albicans* for 2.5 h, and investigated the permeability of the phagolysosome, judged by leaching of SRG pre-loaded to the compartment into the cytosol. Both WT and TMEM63A KO macrophages maintained integrity of the phagolysosomes containing the yeast-locked (YL) *C. albicans*. However, when phagosomes contained Δ*ece1 C. albicans,* a strain that does not produce candidalysin but can undergo hyphal growth, we found that the loss of TMEM63A caused a significant fraction of the phagosomes to undergo rupture (**Fig 3P-Q**). This result is consistent with the notion that TMEM63A confers protection of the lysosomal membrane only upon mechanical challenge.

### Ca^2+^ is not required for the protective effects of TMEM63A

There are several putative mechanisms by which TMEM63A, a non-selective cation channel known to transport calcium, sodium, and potassium (Li et al., 2024), could protect lysosomes from damage. First, TMEM63A could outwardly transport Ca^2+^ to exert protective effects, for example, by recruiting ALG-2 (Chen et al., 2024). This was tested by removing Ca^2+^ from the lysosomes (and the cells) entirely: in the presence of Ca^2+^-free medium, cells were treated with thapsigargin to inhibit the SERCA pump and release Ca^2+^-stores from the ER (**Fig 4A**), reportedly required to supply lysosomes with Ca^2+^ (Calvo et al., 2025; Garrity et al., 2016). We tested if Ca^2+^ was depleted from the lysosomes in this condition by treating the cells with high concentrations of LLOMe to rupture the lysosomes. Here, we did not observe increases in GCaMP6S (**Fig 4B**), a Ca^2+^ biosensor expressed in the cytosol, indicating that the lysosomes were indeed depleted of Ca^2+^ under these conditions. We then treated such cells with concentrations of GPN and LLOMe that caused lysosomal rupture only in the absence of TMEM63A. In lysosomes devoid of Ca^2+^, we did not observe any loss in the protective effects of TMEM63A (**Fig 4C-E**). This would suggest that Ca^2+^ is not relevant cation to confer TMEM63A lysosome protection.

**Figure 4.**
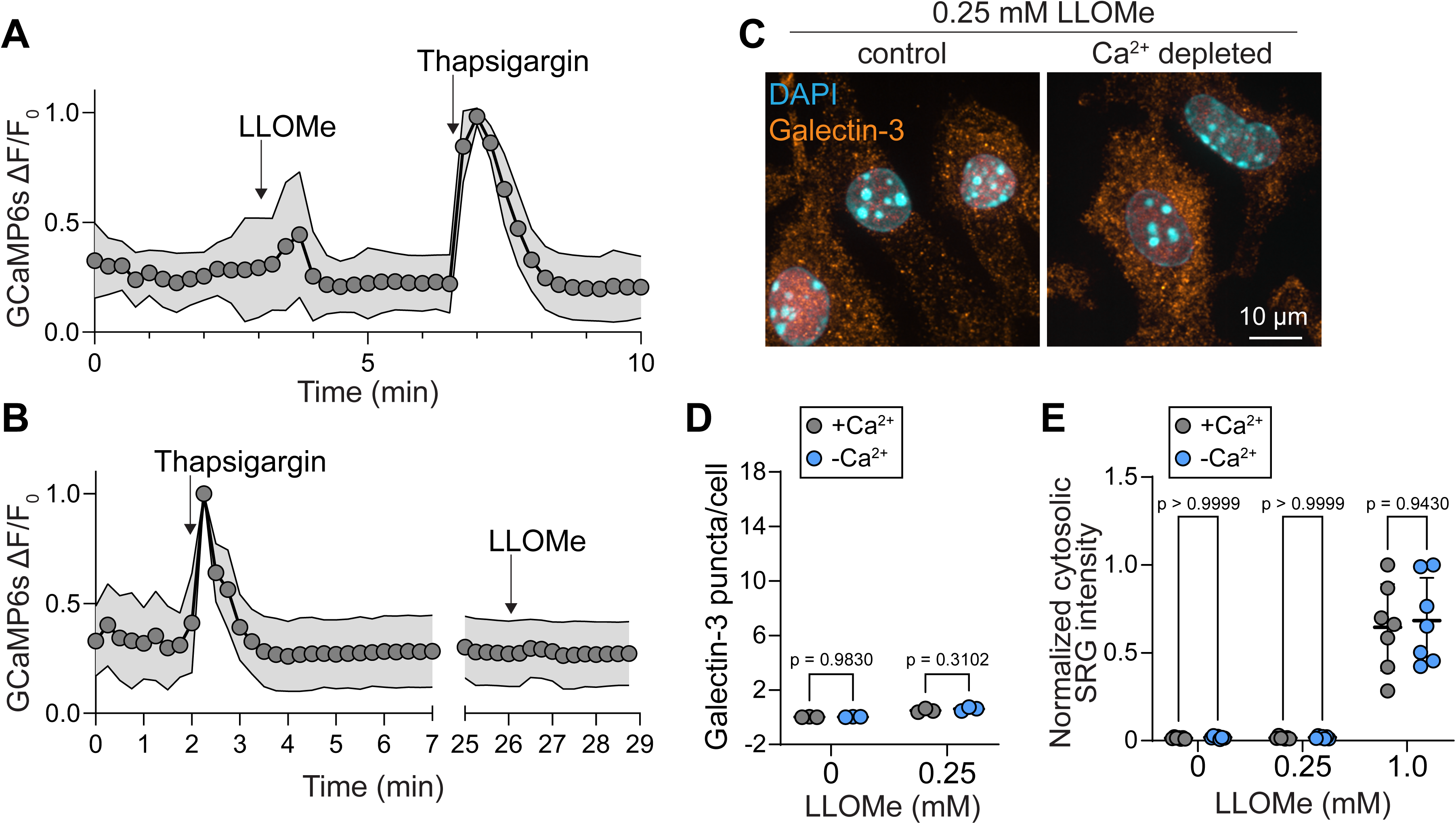
Ca^2+^ efflux is not required for the protective mechanism conferred by TMEM63A. A-B) GCaMP6s recordings from 20-30 WT RAW 264.7 cells. The change in fluorescence (ΔF) was normalized to the highest value for each cell. n = 3. **C-E)** WT BMDM depleted of Ca^2+^ and challenged with LLOMe and quantified as in Figure 1. n = 3, scale bars 10 µm.

### A H^+^-leak and the CASM pathway are not required for the protective effects of TMEM63A

In addition to Ca^2+^ flux, non-selective cation channels like TMEM63A could also result in a H^+^-leak from lysosomes, which could in turn activate lysosome repair pathways. To determine if TMEM63A could transport H^+^, we treated cells with 30 mM of sucrose for 1 h which caused swelling and increased tension as judged by the Lyso-Flipper probe (**S Fig 3A**). We noted that the lysosomes in TMEM63A KO cells in this condition had a modestly more acidic pH (**S Fig 3B**). To monitor the leak of H^+^ in the tensed but intact lysosomes, we inhibited the activity of the V-ATPase with concanamycin A. The rate of lysosomal alkalinization, indicative of the ongoing H^+^ leak, was subtly decreased in the TMEM63A KO cells, suggesting that some of the H^+^ leak was attributed to TMEM63A (**S Fig 3B-C**). We therefore investigated the main pathway activated in response to the stabilization of the V-ATPase and increases in its activity, i.e. the recruitment of LC3 to lysosomes via CASM (Durgan and Florey, 2022). We found that LC3 was clearly recruited to lysosomes upon LLOMe treatment but that this occurred in both WT and TMEM63A KO cells equally (**S Fig 3D**). Alternative insults that selectively ruptured TMEM63A KO cells, on the other hand, did not lead to LC3 recruitment to lysosomes. Thus, it would seem unlikely that TMEM63A protects from lysosome damage by mechanisms related to the H^+^-leak. Instead, we noted that complete inhibition of the V-ATPase with concanamycin A rendered the lysosomes more susceptible to damage (**S Fig 3E)**. These results implicate secondary ion gradients, established by the H^+^ gradient, in the mechanoresilience of lysosomes.

### TMEM63A mediates tension relief and regulated volume decrease of lysosomes

If not Ca^2+^ or H^+^ release, TMEM63A could conceivably protect from lysosomal damage by forcing the exit of water from lysosomes to relieve high membrane tension. This could be achieved by the outward transport of alkali cations, major osmoticants of biological fluids including for lysosomes which contain high luminal Na^+^ (Wang et al., 2012). Volume regulation of lysosomes has indeed been shown to be regulated by VRAC and its transport of Cl^-^, notably under conditions of osmotic stress (Li et al., 2020), which would require the parallel transport of cations to maintain electroneutrality. This seemed like a reasonable possibility since increases in hydrostatic pressure alone caused elevations in endomembrane tension (**Fig 5A**) and led to the loss of luminal Na^+^ as judged by Natrium Green, a pH insensitive dye of the same size as SRG, precluding the possibility that it would be lost from the lumen (**Fig 5B**). The same treatment with hypotonic solutions caused a rapid loss of lysosome tubulation, which was restored over time in control cells (**Fig 5E**). TMEM63A was in fact required for lysosomes to withstand high osmotic pressure: the deletion of TMEM63A caused pronounced loss of lysosome integrity and rupture upon challenging the cells with hypotonic solutions (**Fig 5C-E**). As was also shown for responses to LLOMe, the re-expression of human TMEM63A in TMEM63A KO cells rescued the lysosomal rupture upon hypotonic shock, but loss-of-function variants did not (**S Fig 3G**).

**Figure 5.**
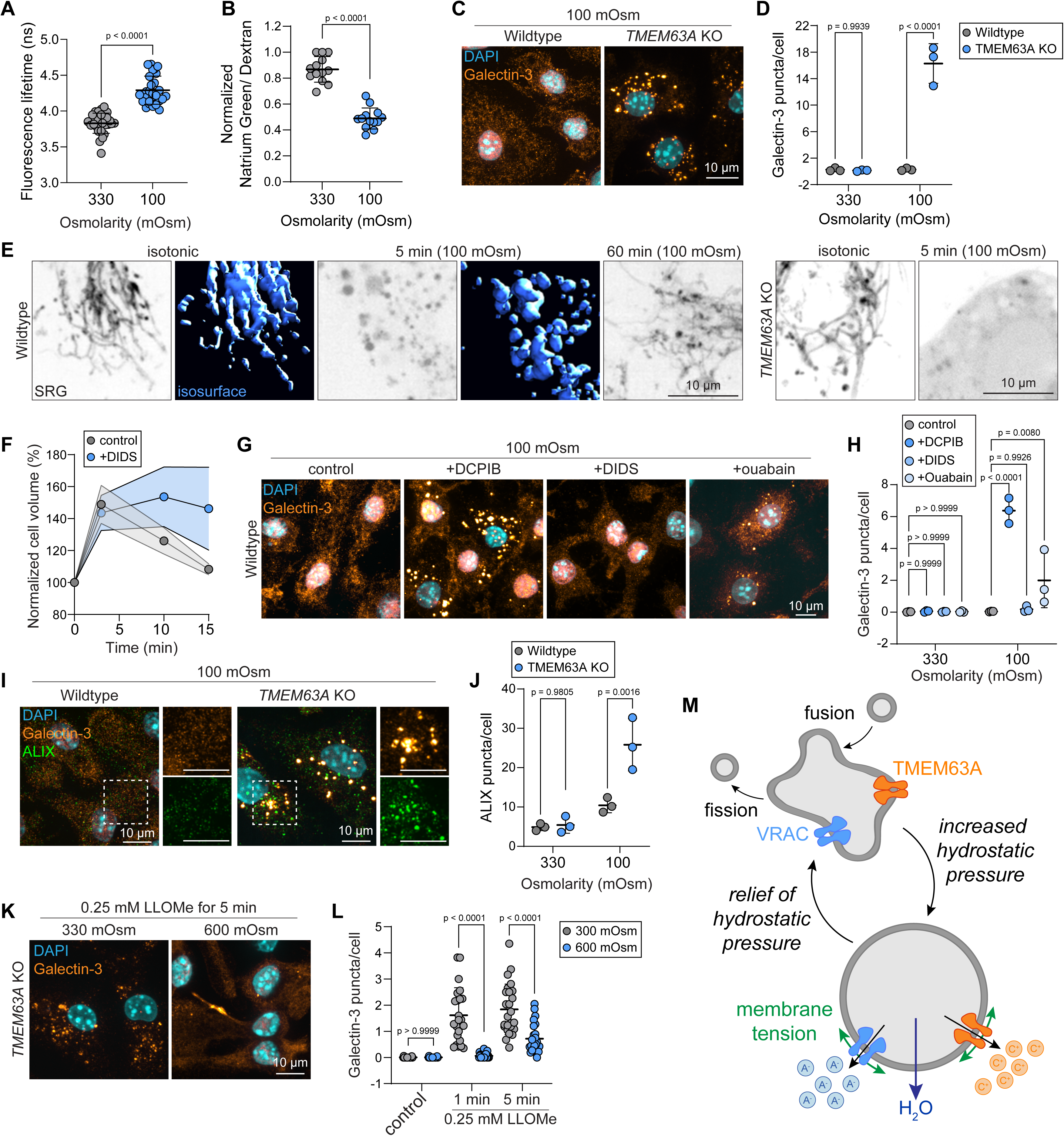
TMEM63A relieves high membrane tension to protect lysosomes from rupture. **A)** Lifetime fluorescence of LysoFlipper WT RAW264.7 cells before and 5 min after hypotonic shock. n = 3. **B)** Relative [Na^+^]_lyso_ of WT BMDM determined using Natrium Green, normalized to 10 kDa dextran, both pulsed for 16 h and chased for 1 h. All data points represent a field containing 4-8 cells. n = 3. **C-D)** Lysosome damage determined for WT and TMEM63A KO BMDM upon 10 min exposure to hypotonic (100 mOsm) solutions. n = 3. **E)** SRG loaded WT BMDM lysosomes undergoing regulatory volume decrease and return to tubulation versus rupture in TMEM63A KO BMDM. *xy* images shown in black and white. Surface rendering of *xyz* images (blue) illustrate tubular versus spherical shapes of the lysosomes. **F)** Cell volumes percentage of WT RAW264.7 cells ± 250 µM DIDS in hypotonic solutions for 3, 10, and 15 min. Normalized to 0 min measurement. n=3. **G-H)** Lysosome damage determined for WT BMDM treated with 100 µM DCPIB for 5 min, 250 µM DIDS for 5min, or 2 mM ouabain for 4 h prior to 10 min exposure to hypotonic (100 mOsm) solutions. n = 3. **I-J)** ALIX recruitment determined for WT and TMEM63A KO BMDM upon 10 min exposure to hypotonic (100 mOsm) solutions. n = 3. **K-L)** Shifting TMEM63A KO BMDM to hypertonic (600 mOsm) solutions at the time of LLOMe delivery. Each data point represents a field containing 4-10 cells. n = 3, scale bars 10 µm. **M)** Model.

There is a very small preference for Na^+^ over K^+^ transport by TMEM63A (Li et al., 2024). Outward transport of Na^+^ would therefore be balanced by the inward transport of K^+^ without any clear net loss of osmolytes if the channel were working in isolation. Instead, the parallel transport of Cl^-^ together with the inside positive membrane potential of the lysosome (Saminathan et al., 2021) would, in theory, result in the directional and electroneutral exit of cations and anions and oblige the exit of water. We therefore opted to inhibit lysosome-resident VRAC using the membrane-permeable inhibitor DCPIB; as a control, we used the membrane-impermeable VRAC inhibitor DIDS which prevented regulatory volume decrease (RVD) as expected (**Fig 5F**). Convincingly, we found that the inhibition of lyso-VRAC with DCPIB but not surface VRAC with DIDS, rendered lysosomes highly susceptible to damage upon hypotonic shock (**Fig 5G-H**). By comparison, when the lysosome-to-cytosol gradient of Na^+^ was blunted by inhibiting the Na^+^/K^+^ pump with ouabain for extended periods, we did not observe damage susceptibility to the same extent, suggesting that either Na^+^ or K^+^ transport suffices for the protective effects of TMEM63A (**Fig 5G-H**). Interestingly, ALIX was only recruited to damaged lysosomes in TMEM63 KO cells and not in WT lysosomes exposed to hypotonic solutions indicating that lysosome repair pathways involving ESCRT were not invoked in this response unless TMEM63A was absent (**Fig 5I-J**).

We then sought to forcibly regulate the volume and resultant tension of lysosomes in the TMEM63A KO macrophages. This was achieved by shifting the osmolarity of the medium upward in order to drive the exit of water from the cells and from the lysosomes at the time of the addition of lysosomotropic agents. These results were revealing as the hypertonic solutions strongly reverted the rupture of lysosomes in the TMEM63A KO macrophages as judged by decreased galectin-3 puncta and decreased release of lysosome contents into the cytosol (**Fig 5K-L**). Taken together, these results implicate a model for TMEM63A in sensing and relieving high tension on lysosomes (**Fig 5M**).

## Discussion

Threats to lysosomal integrity can increase tension on the membrane that delimits the organelle, an effect that causes its rupture if left unregulated. To offset sudden increases in tension, we propose that TMEM63A decreases organelle volume, reduces hydrostatic pressure, and effectively relieves high membrane tension. This could provide membrane slack and time for the organelle to obtain additional membrane via fusion with other lysosomes or lipid transport from the ER. Supporting this model, lysosomes experiencing sudden osmotic stress undergo a period of growth/expansion before undergoing secondary volume loss. These stages are not observed in the absence of TMEM63A because the stressed organelles undergo rapid lysis. The slow accumulation of solutes, on the other hand, proceeds without damage and without the need for TMEM63A. In such cases, fusion and/or the addition of ER lipids clearly suffices to maintain organelle integrity and occurs independently of TMEM63A.

The OSCA/TMEM63 family of mechanosensitive channels is highly conserved through evolution, members of which can be found in plants, including on their vacuoles. Osmotic stresses in plants –incurred upon injuries to the cell wall– lead to the mechanical gating these channels, presumably enabling the free passage of cations (Cao et al., 2020; Miao et al., 2022; Yuan et al., 2014). The expression of OSCA channels ultimately preserves vacuolar integrity upon cell wall damage and supports the viability of plants, which is akin to our model for TMEM63A on lysosomes.

While calcium would stand to be released through TMEM63A and this could increase membrane fusion, we did not find there to be a role in whole cell calcium for preserving lysosome integrity in very acute responses to lysosomotropic agents. Rather than using BAPTA-AM which is cleaved by cytosolic esterases and accumulates in cells together with its toxic byproducts, formaldehyde and acetic acid, we instead depleted cells of their Ca^2+^ stores entirely, including those in lysosomes. Calcium is a critical mediator of plasma membrane and endomembrane repair, notably by its activation of the ESCRT machinery (Vietri et al., 2020). We found that the recruitment of ALIX, a protein that recruits the ESCRT-III to endomembranes, was unaffected in TMEM63A KO cells, suggesting that Ca^2+^ release or ESCRT-III recruitment are not responsible for the immediate protective effects of TMEM63A. A steady leak of cations or the decrease in membrane tension facilitated by TMEM63A and its role in ESCRT recruitment and activity before rupture are nevertheless possible.

A critical, outstanding question is how lysosomes gain surface to respond to insults. One dramatic example occurs in contexts in which phagosomes need to expand to accommodate rapidly growing pathogens harboured in their lumen. In such instances, lysosomes undergo substantive fusion with the phagosome which begins to consume the entire population of LAMP1-positive compartments (Westman et al., 2020). Such homotypic fusion is also initiated when PIKfyve is inhibited leading to vacuolation (Choy et al., 2018). An alternative or perhaps additional mechanism comes from lipid delivery from the ER via lipid-binding proteins, e.g. ATG2, VPS13C, and PDZD8. Lipid transport would need to also be coupled to scrambling. In that regard, it is interesting to note that TMEM63B can function as a lipid scramblase (Miyata et al., 2025) which also targets to lysosomes (Li et al., 2024). It may be interesting to investigate the potential coupling between TMEM63A, B, and the bridge-like lipid transfer proteins.

The role of TMEM63A in protecting lysosomes from damage to their membrane has immediate implications in inflammation and antigen cross-presentation (Canton et al., 2021). Since the channel protects from lysosomal damage, inactivation of the channel may increase inflammatory processes and the release of tumor-associated antigens. Other mechanisms of lysosomal resilience are surely also in place including biophysical mechanisms that protect the bilayer like the lysosomal glycocalyx, membrane stabilization provided by ESCRT machinery, and potentially yet to be determined ion transport pathways (Scott et al., 2025). This study establishes a framework whereby lysosomal resilience is considered greater than first appreciated, which would need be taken into account when understanding the lost protection in pathology associated with lysosomal damage.

## Methods

### Cell isolation, culture, and transfection

Primary murine macrophages were derived from the marrow of femoral bones from 6–8-weekold C57Bl/6 wild-type, LysM-Cre_floxed;floxed TMEM63a (*Tmem63a*^−/−^) and littermate LysMCre_ WT;floxed TMEM63a (wildtype) control mice. 2-5 littermates per cage were housed in ventilated cages under standard housing conditions (40% humidity, 22°C, 12 h reversed light/dark cycle) with free access to chow diet. All procedures were approved by the Animal Care Committee at the Hospital for Sick Children and were performed in accordance with regulations from the Animals for Research Act of Ontario and the Canadian Council on Animal Care. Mice were kept at the Animal Facility in the Peter Gilgan Center for Research and Learning.

BMDM were harvested by centrifugation and grown in Dulbecco’s modified Eagle medium (DMEM) with L-glutamine-containing 10% heat-inactivated foetal calf serum (FBS) and 100 U ml^−1^ penicillin and 100 μg ml^−1^ streptomycin. BMDM were plated at 1 × 10^6^ cells ml^−1^ and differentiated using 10% L929-conditioned medium for 5 days at 37 °C with 5% CO_2_ on Petri dishes. Cells were lifted using 5 mM EDTA in PBS, pH 7.4, and seeded onto coverslips.

RAW264.7 were grown and passaged in DMEM + 10% FBS at 37 °C with 5% CO_2_. RAW264.7 TMEM63A KO cells were generated as previously described (Cai et al., 2024). Transfection of RAW264.7 cells using FuGeneHD was done according to the manufacturer’s protocol.

### Sulforhodamine G imaging

Cells seeded onto coverslips were pulsed with 5 mg/mL sulforhodamine G (Molecular probes, S-6957) up to overnight and chased for >1 h before the addition of LLOMe, GPN, alum, sucrose, pathogens or hypotonic solutions. Cells were imaged live in HBSS (Wisent, 311-512 CL). The mean fluorescence of cytosolic SRG signal of each cell was determined using three regions of interest, subtracting background signal, and normalized to the highest value (i.e. lysosomal) for each experiment.

### Live cell imaging

A 63×1.35 numerical aperture oil immersion objective using an Olympus IX81 Quorum spinning disc confocal microscope was used to image. The microscope was equipped with a CSU-X1-A Yokogawa spinning-disc unit and a Hamamatsu C9100-13 EM-CCD camera controlled by Volocity 6.1.2 software (PerkinElmer). Fluorescence images shown as volume-render projections belong to Z-stacks acquired at 0.4 μm.

### Immunofluorescence

RAW264.7 cells and BMDM were seeded on 18 mm coverslips in 12-well tissue culture plates 16 h prior to experimentation. Cells were fixed and permeabilized with ice cold 100% MeOH for 5 min at -20 °C and blocked in 1% BSA for 1 h. Subsequently, the cells were incubated with the indicated primary and secondary antibodies in 1% BSA for 1.5 h at room temperature. All primary antibodies and Alexa488-, Cy3- or Cy5-conjugated secondary antibodies (Jackson ImmunoResearch) diluted to 1:1000 in blocking solution. Three washes with 1X PBS were performed after each incubation. Slides were mounted using mounting media containing DAPI (Invitrogen, P36931).

### Lyso Flipper TR and FLIM

RAW264.7 cells were seeded on 18 mm coverslips in 12-well tissue culture plates 16 h prior to experimentation. Cells were incubated with 2 μm Lyso Flipper TR (Cytoskeleton, CY-SC022) at 37°C for 10 min in serum-free DMEM before washing 3 times with PBS and imaging in HBSS. Fluorescence-lifetime imaging microscopy (FLIM) was acquired using the Leica Stellaris 8 FALCON w/STED. The Leica LAS X software (Stellaris) was used for□analysis.

### Quantitative RT-PCR (RT-qPCR)

RAW264.7, BMDM and mouse embryonic fibroblasts were plated on 6-well tissue culture plates and harvested 16 h after. Cellular RNA was extracted using the GeneJET RNA purification kit (Thermo Fisher Sceintific). Equal RNA from each condition was used to generate cDNA with the Superscript VILO cDNA Synthesis Kit (Thermo Fisher Scientific). Quantitative real-time PCR was then performed with TaqMan probes specific to each assayed gene on a QuantStudio 3 Real-Time PCR System (Applied Biosystems, Thermo Fisher Scientific) in 96-well plates. TaqMan probes targeting mouse genes Tmem63a (Mm00522653_m1; no. 4331182), Tmem63b (Mm01269695_m1; no. 4351372), and Tmem63c (Mm00553595_m1; no. 4331182) were from Thermo Fisher (no. 4448892). Readouts for target and reference (Abt1) genes were performed in duplicate. Relative gene expression was then calculated based on the 2^−ΔΔCt^□method by comparing the relative quantification of the control gene to each target gene.

### Ratiometric pH determinations

Cells seeded on coverslips were pulsed with 25□µg□/mL of 10 kDa Oregon green dextran (Thermo Fisher Scientific, D7170) overnight and chased for >1 h before measurement. The cells were excited at 490 and 440 nm wavelengths in HBSS buffer, followed by incubation of pH standard solutions with pH values at 4.5, 5.5, 6.5 and 7.5 to obtain a standard pH curve. The solutions were made with 143 mM KCl, 5 mM glucose, 1 mM MgCl_2_ and 1 mM CaCl_2_ buffered with 20 mM acetic acid (pH 4.5), 20 mM MES (pH 5.5 and 6.5) or 20 mM HEPES (pH 7.5). A concentration of 10 μM nigericin (Cayman chemical, 11437) and 5 μM monensin (Sigma-Aldrich, 475895) were added to each pH buffer before use. The curve was graphed from the background subtracted values for the ratiometric fluorescence (490/440 nm). The corresponding baseline pH was calculated from the standard curve.

Imaging was performed on a microscopy system that consists of a microscope (Axio Observer, Zeiss), a cooled CCD camera (Evolve 512, Photometrics), and a fluorescent light source (HXP 120V, Zeiss). Ratiometric imaging was performed with two filter cubes: a 470/40 nm and a 436/20 nm. The microscope is operated using ZEN 2 blue edition software (Zeiss), and images were acquired with a 63×/1.4 NA oil objective (Zeiss).

### DQ-BSA assay

BMDM were seeded on 18 mm coverslips in 12-well tissue culture plates 16 h prior to imaging. Cells were loaded with 10 µg/mL DQ-BSA Red (Invitrogen, D12051) for 1 h and imaged in HBSS using confocal microscopy.

### Phagocytosis assay

BMDM were seeded on 18 mm coverslips in 12-well tissue culture plates 16 h prior to imaging. Cells were then incubated with human IgG-opsonized 4.98 μm silica beads (Bangs Laboratories Inc., SS05003) for 30 min. Cultures were then fixed with 3% PFA for 20 min and incubated with fluorescent anti-human secondary antibodies for 1 h at room temperature to visualize beads that were not ingested by the macrophages. Cells were then permeabilized with 0.1% Triton-x for 5 min and incubated with a different fluorescent anti-human secondary antibody to visualize internalized beads for 1 h. Actin was stained with phalloidin to delineate the macrophages.

### PI assay

RAW264.7 cells were grown to 80% confluency in 6-well plates and challenged with the indicated concentrations of LLOMe for 30 min. After 25 min, 0.3 µg/mL of PI (Invitrogen, P1304MP) was added for 5 min. Three images of different fields of view of each well were taken in both RFP and brightfield setting of an EVOS microscope (Thermo Fisher Scientific). Quantification of PI positivity was done in ImageJ.

### Candida albicans assay

*C. albicans* cultures were grown at 30°C overnight in liquid yeast peptone dextrose (YPD) medium (Bioshop). Overnight *C. albicans* cultures were diluted to OD_600nm_□∼0.1 in YPD media and grown at 30°C until OD_600nm_□reached 1.0. Prior to infection, *C. albicans* were opsonized with human serum for 60 min at room temperature with rotating. The subculture was diluted into DMEM containing 10% FBS. Using a 270-gauge needle, *C. albicans* aggregates were dispersed. RAW264.7 wildtype and TMEM63A KO cells were seeded on 18 mm coverslips in 12-well plates and were pulsed with SRG overnight. For infections, the medium was removed from the wells and replaced with 1 mL of *C. albicans* containing DMEM. Plates were centrifuged at 1500 rpm for 1 min, then incubated at 37 °C and 5% CO_2_□for 2.5 h. Cells were then imaged live in HBSS.

### Ca^2+^ imaging

RAW264.7 cells transfected with GCaMP6s were washed with PBS three times, followed by incubation in Ca^2+^ free Tyrode’s buffer (10 mM HEPES, 10 mM D-glucose, 5 mM KCl, 140 mM NaCl, 1 mM MgCl_2_, 1 mM EGTA) for 5 min. Cells were recorded every 15 s in Ca^2+^ free Tyrode’s buffer and were treated with either 2 µM thapsigargin or 3 mM of LLOMe. Samples were imaged by spinning disc confocal microscopy using a 25x objective. The change in GCaMP6s fluorescence was quantified by normalizing to the highest value for each cell.

### Determination of H^+^□leak

Wildtype and□*TMEM63A□*KO RAW264.7 cells seeded on 18 mm coverslips in 12-well tissue culture plates were incubated with 10 kDa Oregon Green dextran overnight and chased for >1 h before measurement. The pH of the lysosomes was measured ratiometrically over 18 min, and cells were acutely treated with 500 nM concanamycin A (Cayman Chemical, 11050) at 4 min time point.

### In situ measurement of lysosomal cations

BMDM were seeded on 18 mm coverslips in 12-well tissue culture plates and were loaded with 25□µg□/mL of 10 kDa Alexa fluor-647 dextran (Thermo Fisher Scientific, D22914) and 5 mg/mL ING-2 TMA+ salt (ION Biosciences) overnight and chased for >1 h before measurement. Cells were imaged in HBSS using live confocal microscopy.

### Cell volume measurement

RAW264.7 cells were grown to 80% confluency in 6-well plates. After, they were lifted mechanically. 0.5 mL of collected cell suspension of each condition was diluted with 3 mL Isoton II Diluent 531 (Beckman Coulter) and 6 mL distilled water immediately before analysis with a Coulter counter (Beckman Coulter Life Sciences, Multisizer 4). The median cell diameter for >10000 cells ranging from 5 - 25 µm was measured at 0, 3, 10, and 15 min. The diameter was converted to volume by assuming that the suspended cells were spherical. Values were normalized to the 0 min time point.

### Confocal microscopy

Confocal microscopy was performed on a spinning-disk system (Quorum Technologies Inc.), consisting of a microscope (Axiovert 200 M, Zeiss), cooled CCD camera (ORCA-Fusion BT, Hamamatsu), five-line laser module (Spectral Applied Research) with 405-, 443-, 491-, 561-, and 655-nm lines, and a filter wheel (MAC5000, Ludl). The microscope is operated using Volocity v6.3 software (Perkin Elmer). Images were acquired with a 63×/1.4 NA oil objective (Zeiss).

### Plasmids

TMEM63A-OFP was purchased from Sino Biological (MG51287-ACR). G168E and G567S were generated with site-directed mutagenesis using Q5 High-Fidelity DNA Polymerase (New England BioLabs, M0494). LAMP1-GFP was a gift from Ron Vale (Addgene plasmid # 16290). EEA1-GFP was a gift from Silvia Corvera (Addgene plasmid # 42307). GCaMP6s was a gift from Douglas Kim & GENIE Project (Addgene plasmid # 40753).

### Reagents

L-Leucyl-L-Leucine methyl ester (Sigma-Aldrich, L7393), Gly-phe-β-napthylamide (Cayman Chemical, 14634), alum hydroxide (Invivogen, 21645-51-2), and sucrose (Millipore Sigma, 57-501) were used at indicated concentrations. Anti-galectin-3 (abcam, ab76245) and ALIX (BioLegend, 634502) were used at 1:1000 (v/v) for immunofluorescence. LysoTracker Red DND-99 (Thermo Fisher Scientific, L7528) was used at 1 µM. Primary antibodies against LAMP1 (DSHB, 1D4B), Cathepsin C (Santa Cruz, sc-74590) and GAPDH (Santa Cruz, 365062) were used at 1:1000 (v/v) for western blotting. Thapsigargin (Calbiochem, 586005), ouabain (Sigma Aldrich, O3152), DCPIB (Tocris, 1540), and DIDS (Sigma-Aldrich, D3514) were used at 2 µM, 2 mM, 100 µM, and 250 µM respectively.

## Acknowledgements

S.A.F. is the recipient of a Canada Research Chair and supported by grants PJT-169180 from the Canadian Institutes of Health Research (CIHR) and RGPIN-2022-04485 from the Natural Sciences and Engineering Research Council of Canada (NSERC). All graphics were created with BioRender.com.

## References

Cai, R., O. Scott, G. Ye, T. Le, E. Saran, W. Kwon, S. Inpanathan, B.A. Sayed, R.J. Botelho, A. Saric, S. Uderhardt, and S.A. Freeman. 2024. Pressure sensing of lysosomes enables control of TFEB responses in macrophages. Nat Cell Biol. 26:1247–1260.

Calvo, B., P. Torres-Vidal, A. Delrio-Lorenzo, C. Rodriguez, F.J. Aulestia, J. Rojo-Ruiz, B. Callejo, B.M. McVeigh, M. Keller, C. Grimm, V. Oorschot, V. Moiseenkova-Bell, D.I. Yule, J. Garcia-Sancho, S. Patel, and M.T. Alonso. 2025. Direct measurements of luminal Ca2+ with endo-lysosomal GFP-aequorin reveal functional IP3 receptors. J Cell Biol. 224.

Canton, J., H. Blees, C.M. Henry, M.D. Buck, O. Schulz, N.C. Rogers, E. Childs, S. Zelenay, H. Rhys, M.C. Domart, L. Collinson, A. Alloatti, C.J. Ellison, S. Amigorena, V. Papayannopoulos, D.C. Thomas, F. Randow, and C. Reis e Sousa. 2021. Publisher Correction: The receptor DNGR-1 signals for phagosomal rupture to promote cross-presentation of dead-cell-associated antigens. Nat Immunol. 22:391.

Cao, L., P. Zhang, X. Lu, G. Wang, Z. Wang, Q. Zhang, X. Zhang, X. Wei, F. Mei, L. Wei, and T. Wang. 2020. Systematic Analysis of the Maize OSCA Genes Revealing ZmOSCA Family Members Involved in Osmotic Stress and ZmOSCA2.4 Confers Enhanced Drought Tolerance in Transgenic Arabidopsis. Int J Mol Sci. 21.

Chen, W., M.M. Motsinger, J. Li, K.P. Bohannon, and P.I. Hanson. 2024. Ca(2+)-sensor ALG-2 engages ESCRTs to enhance lysosomal membrane resilience to osmotic stress. Proc Natl Acad Sci U S A. 121:e2318412121.

Choy, C.H., G. Saffi, M.A. Gray, C. Wallace, R.M. Dayam, Z.A. Ou, G. Lenk, R. Puertollano, S.C. Watkins, and R.J. Botelho. 2018. Lysosome enlargement during inhibition of the lipid kinase PIKfyve proceeds through lysosome coalescence. J Cell Sci. 131.

Colom, A., E. Derivery, S. Soleimanpour, C. Tomba, M.D. Molin, N. Sakai, M. Gonzalez-Gaitan, S. Matile, and A. Roux. 2018. A fluorescent membrane tension probe. Nat Chem. 10:1118–1125.

Duewell, P., H. Kono, K.J. Rayner, C.M. Sirois, G. Vladimer, F.G. Bauernfeind, G.S. Abela, L. Franchi, G. Nunez, M. Schnurr, T. Espevik, E. Lien, K.A. Fitzgerald, K.L. Rock, K.J. Moore, S.D. Wright, V. Hornung, and E. Latz. 2010. NLRP3 inflammasomes are required for atherogenesis and activated by cholesterol crystals. Nature. 464:1357–1361.

Durgan, J., and O. Florey. 2022. Many roads lead to CASM: Diverse stimuli of noncanonical autophagy share a unifying molecular mechanism. Sci Adv. 8:eabo1274.

Evans, E.A., R. Waugh, and L. Melnik. 1976. Elastic area compressibility modulus of red cell membrane. Biophys J. 16:585–595.

Filimonenko, M., S. Stuffers, C. Raiborg, A. Yamamoto, L. Malerod, E.M. Fisher, A. Isaacs, A. Brech, H. Stenmark, and A. Simonsen. 2007. Functional multivesicular bodies are required for autophagic clearance of protein aggregates associated with neurodegenerative disease. J Cell Biol. 179:485–500.

Gaillard, J.L., P. Berche, J. Mounier, S. Richard, and P. Sansonetti. 1987. In vitro model of penetration and intracellular growth of Listeria monocytogenes in the human enterocyte-like cell line Caco-2. Infect Immun. 55:2822–2829.

Garrity, A.G., W. Wang, C.M. Collier, S.A. Levey, Q. Gao, and H. Xu. 2016. The endoplasmic reticulum, not the pH gradient, drives calcium refilling of lysosomes. Elife. 5.

Hornung, V., F. Bauernfeind, A. Halle, E.O. Samstad, H. Kono, K.L. Rock, K.A. Fitzgerald, and E. Latz. 2008. Silica crystals and aluminum salts activate the NALP3 inflammasome through phagosomal destabilization. Nat Immunol. 9:847–856.

Koslov, M.M., and V.S. Markin. 1984. A theory of osmotic lysis of lipid vesicles. J Theor Biol. 109:17–39.

Li, K., Y. Guo, Y. Wang, R. Zhu, W. Chen, T. Cheng, X. Zhang, Y. Jia, T. Liu, W. Zhang, L.Y. Jan, and Y.N. Jan. 2024. Drosophila TMEM63 and mouse TMEM63A are lysosomal mechanosensory ion channels. Nat Cell Biol. 26:393–403.

Li, P., M. Hu, C. Wang, X. Feng, Z. Zhao, Y. Yang, N. Sahoo, M. Gu, Y. Yang, S. Xiao, R. Sah, T.L. Cover, J. Chou, R. Geha, F. Benavides, R.I. Hume, and H. Xu. 2020. LRRC8 family proteins within lysosomes regulate cellular osmoregulation and enhance cell survival to multiple physiological stresses. Proc Natl Acad Sci U S A. 117:29155–29165.

Miao, S., F. Li, Y. Han, Z. Yao, Z. Xu, X. Chen, J. Liu, Y. Zhang, and A. Wang. 2022. Identification of OSCA gene family in Solanum habrochaites and its function analysis under stress. BMC Genomics. 23:547.

Miyata, Y., K. Takahashi, Y. Lee, C.S. Sultan, R. Kuribayashi, M. Takahashi, K. Hata, T. Bamba, Y. Izumi, K. Liu, T. Uemura, N. Nomura, S. Iwata, S. Nagata, T. Nishizawa, and K. Segawa. 2025. Membrane structure-responsive lipid scrambling by TMEM63B to control plasma membrane lipid distribution. Nat Struct Mol Biol. 32:185–198.

Murthy, S.E., A.E. Dubin, T. Whitwam, S. Jojoa-Cruz, S.M. Cahalan, S.A.R. Mousavi, A.B. Ward, and A. Patapoutian. 2018. OSCA/TMEM63 are an Evolutionarily Conserved Family of Mechanically Activated Ion Channels. Elife. 7.

Niekamp, P., F. Scharte, T. Sokoya, L. Vittadello, Y. Kim, Y. Deng, E. Sudhoff, A. Hilderink, M. Imlau, C.J. Clarke, M. Hensel, C.G. Burd, and J.C.M. Holthuis. 2022. Ca(2+)-activated sphingomyelin scrambling and turnover mediate ESCRT-independent lysosomal repair. Nat Commun. 13:1875.

Saminathan, A., J. Devany, A.T. Veetil, B. Suresh, K.S. Pillai, M. Schwake, and Y. Krishnan. 2021. A DNA-based voltmeter for organelles. Nat Nanotechnol. 16:96–103.

Scott, O., E. Saran, and S.A. Freeman. 2025. The spectrum of lysosomal stress and damage responses: from mechanosensing to inflammation. EMBO Rep. 26:1425–1439.

Skowyra, M.L., P.H. Schlesinger, T.V. Naismith, and P.I. Hanson. 2018. Triggered recruitment of ESCRT machinery promotes endolysosomal repair. Science. 360.

Thiele, D.L., and P.E. Lipsky. 1990. Mechanism of L-leucyl-L-leucine methyl ester-mediated killing of cytotoxic lymphocytes: dependence on a lysosomal thiol protease, dipeptidyl peptidase I, that is enriched in these cells. Proc Natl Acad Sci U S A. 87:83–87.

Vietri, M., M. Radulovic, and H. Stenmark. 2020. The many functions of ESCRTs. Nat Rev Mol Cell Biol. 21:25–42.

Wang, X., P. Xu, A. Bentley-DeSousa, W. Hancock-Cerutti, S. Cai, B.T. Johnson, F. Tonelli, L. Shao, G. Talaia, D.R. Alessi, S.M. Ferguson, and P. De Camilli. 2025. The bridge-like lipid transport protein VPS13C/PARK23 mediates ER-lysosome contacts following lysosome damage. Nat Cell Biol. 27:776–789.

Wang, X., X. Zhang, X.P. Dong, M. Samie, X. Li, X. Cheng, A. Goschka, D. Shen, Y. Zhou, J. Harlow, M.X. Zhu, D.E. Clapham, D. Ren, and H. Xu. 2012. TPC proteins are phosphoinositide-activated sodium-selective ion channels in endosomes and lysosomes. Cell. 151:372–383.

Westman, J., G.F.W. Walpole, L. Kasper, B.Y. Xue, O. Elshafee, B. Hube, and S. Grinstein. 2020. Lysosome Fusion Maintains Phagosome Integrity during Fungal Infection. Cell Host Microbe. 28:798–812 e796.

Yan, H., G. Helman, S.E. Murthy, H. Ji, J. Crawford, T. Kubisiak, S.J. Bent, J. Xiao, R.J. Taft, A. Coombs, Y. Wu, A. Pop, D. Li, L.S. de Vries, Y. Jiang, G.S. Salomons, M.S. van der Knaap, A. Patapoutian, C. Simons, M. Burmeister, J. Wang, and N.I. Wolf. 2019. Heterozygous Variants in the Mechanosensitive Ion Channel TMEM63A Result in Transient Hypomyelination during Infancy. Am J Hum Genet. 105:996–1004.

Yang, H., J. Xun, Y. Li, A. Mondal, B. Lv, S.C. Watkins, L. Shi, and J.X. Tan. 2025. LYVAC/PDZD8 is a lysosomal vacuolator. Science. 389:eadz0972.

Yuan, F., H. Yang, Y. Xue, D. Kong, R. Ye, C. Li, J. Zhang, L. Theprungsirikul, T. Shrift, B. Krichilsky, D.M. Johnson, G.B. Swift, Y. He, J.N. Siedow, and Z.M. Pei. 2014. OSCA1 mediates osmotic-stress-evoked Ca2+ increases vital for osmosensing in Arabidopsis. Nature. 514:367–371.

