## Supplemental Figures for "Mechanoresilience of lysosomes conferred by TMEM63A"

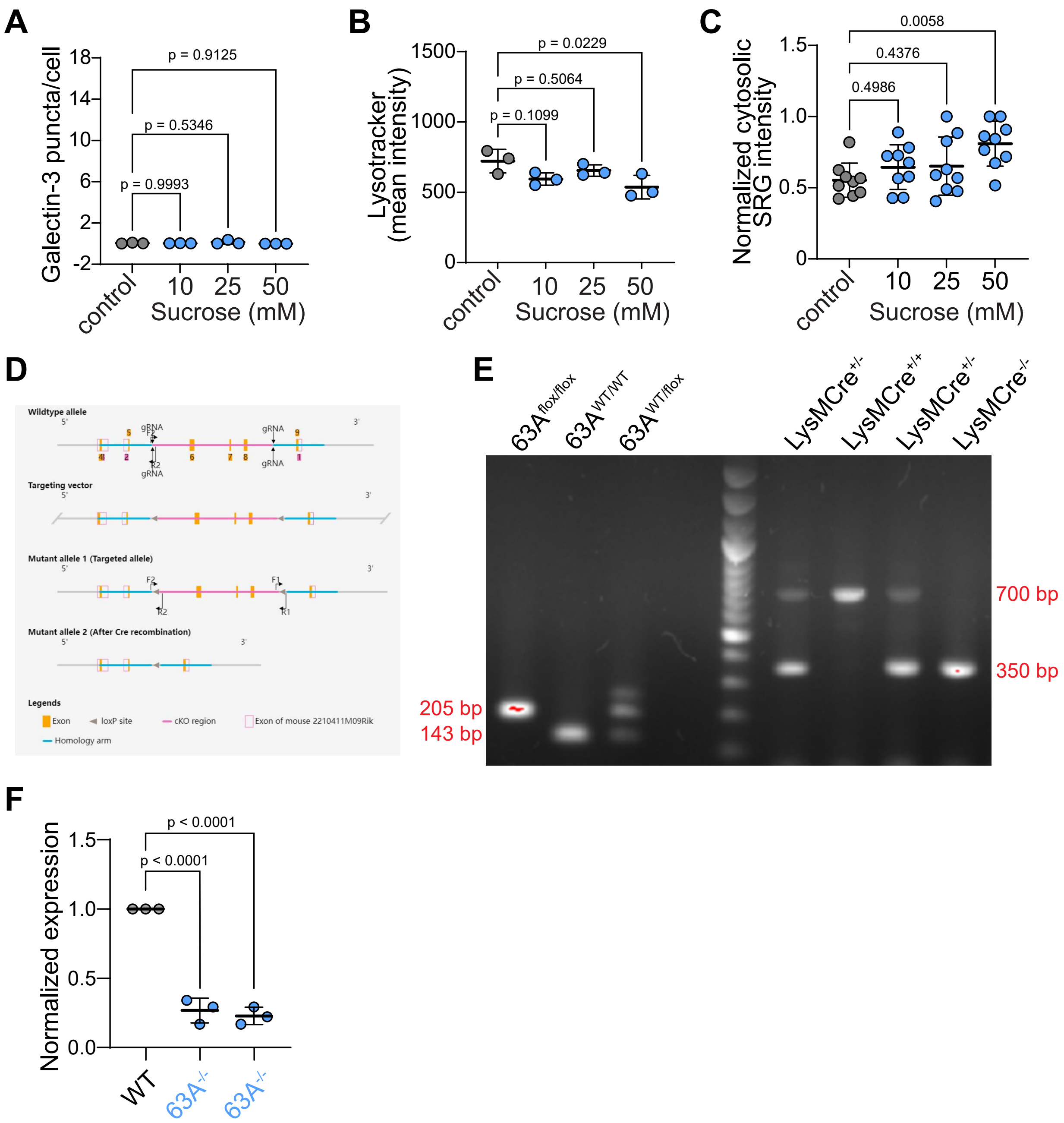


**Supplemental Figure 1**. **A-C)** Primary macrophages with indicated concentrations of sucrose for 16 h. In each case, the damage to lysosomes was determined by fixing and staining for galectin puncta (each dot represents mean of experiment), quantifying the lysotracker intensity per cell (>5 cells per field, 15 fields, each dot represents mean of experiment), or sulforhodamine G (SRG, >5 cells per field, 9 fields, each dot representing a field). n = 3. **D)** Diagram indicating the LoxP sites flanking the 6 to 8 exons of *Tmem63a*. **E)** Genotyping of Tmem63a$^{flox/flox}$ and LysMCre. **F)** qPCR for WT and TMEM63A KO BMDM. n = 3.

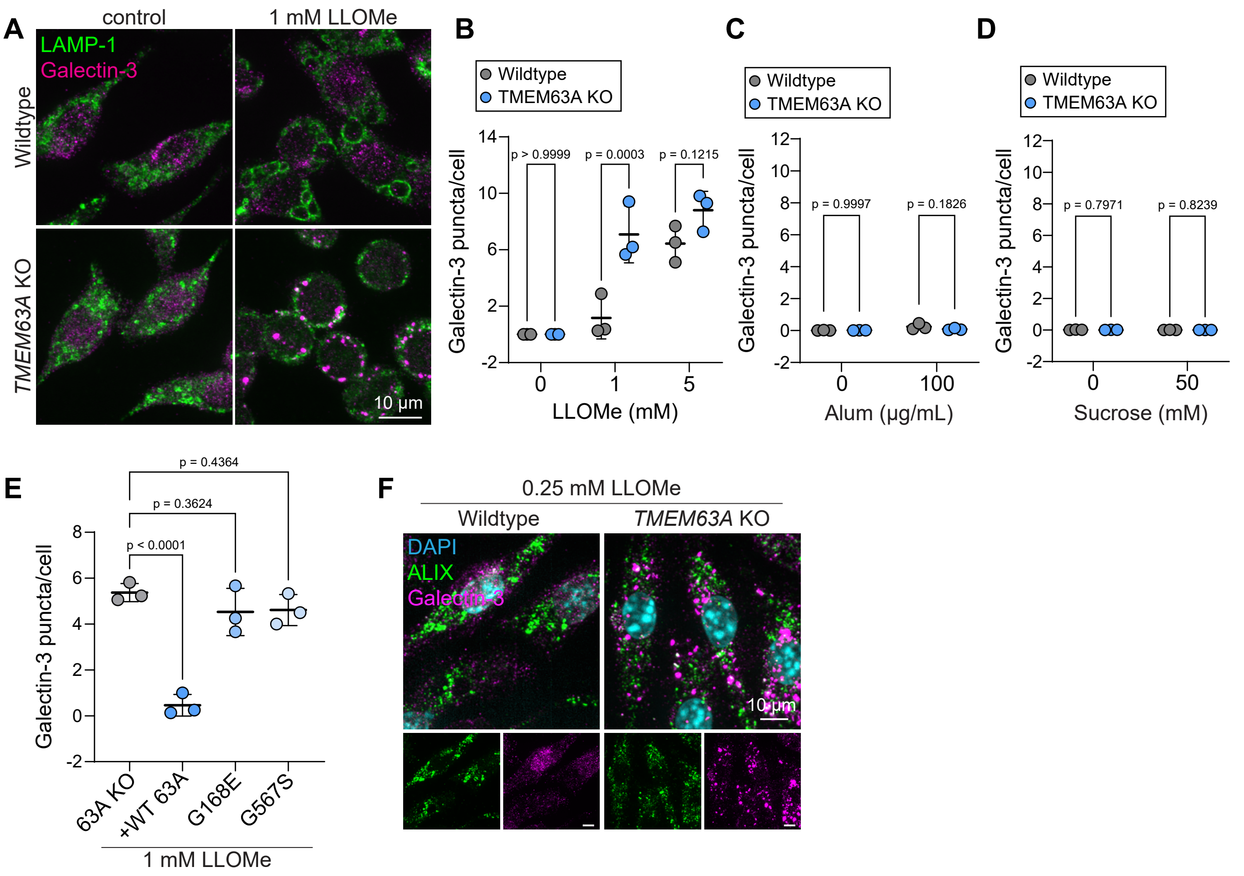


**Supplemental Figure 2**. **A-D)** In all cases, the damage response for WT and TMEM63A KO RAW264.7 cells was determined by staining for galectin-3 puncta along with LAMP-1. **A-B)** LLOMe at indicated concentrations was given to the cells for 20 min. **C)** 100 µg/mL alum was given for 3 h. **D)** 50 mM sucrose was given overnight for 16 h. These experiments were done for multiple TMEM63A KO clones, 3-5 fields per experiment. n=3. **E)** TMEM63A KO RAW264.7 cells re-expressing a TMEM63A-OFP or indicated variants of TMEM63A-OFP and challenged with 1 mM LLOMe for 20 min. Galectin puncta were quantified. n=3. **F)** WT or TMEM63A KO BMDM challenged with 0.25 mM LLOMe for 10 min were immunostained with galectin-3 (magenta) and ALIX (green).

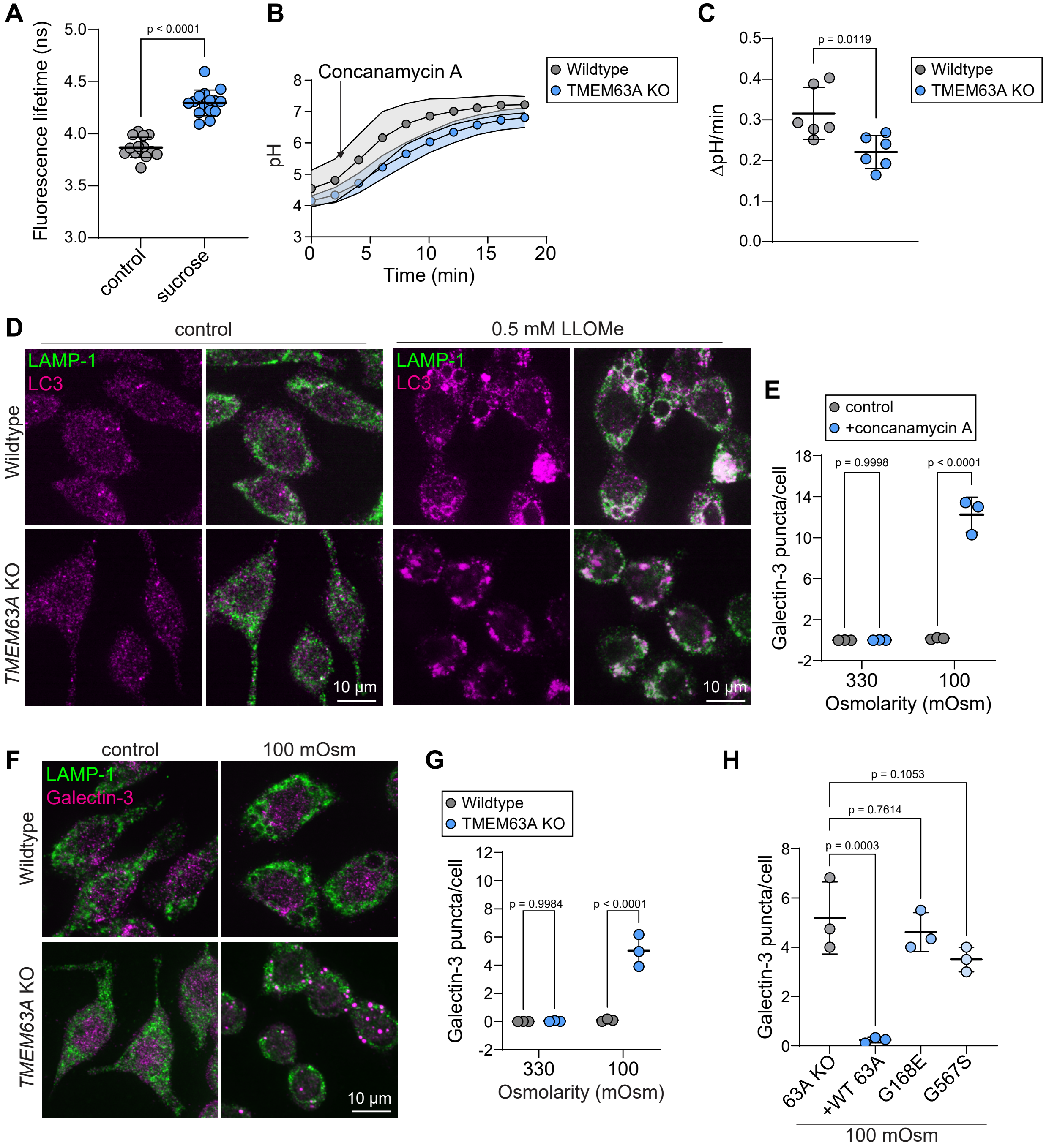


**Supplemental Figure 3**. **A)** Lyso Flipper FLIM in RAW264.7 macrophages incubated with 30 mM sucrose overnight. Each dot represents one field of 4-8 cells. n = 3. **B)** $H^+$-leak determination using ratiometric measurements of Oregon Green 10 kDa dextran after concanamycin addition. n = 3. **C)** Mean instantaneous rates of pH changes from 2 to 8 min. **D)** Examples of WT or TMEM63A KO RAW 264.7 cells challenged with 0.5 mM LLOMe for 15 min. Cell were immunostained with LAMP-1 (green) and LC3 (magenta). **E)** Quantification of galectin-3 puncta with WT BMDM pre-treated with 5 µM of concanamycin A for 30 min prior to hypotonic (100 mOsm) shock. n = 3. **F-G)** WT and TMEM63A KO RAW 264.7 cells were incubated in hypotonic (100 mOsm) solutions for 15 min. Immunostained with galectin-3 (magenta) and LAMP-1 (green), quantified in G. n = 3. **H)** TMEM63A KO RAW264.7 cells re-expressing a OFP version of the channel and challenged with 100 mOsm solution for 15 min. Galectin puncta were quantified (each dot representing mean of experiment). n=3.
